# Phylogenetic pattern of SARS-CoV-2 from COVID-19 patients from Bosnia and Herzegovina: lessons learned to optimize future molecular and epidemiological approaches

**DOI:** 10.1101/2020.06.19.160606

**Authors:** Teufik Goletic, Rijad Konjhodzic, Nihad Fejzic, Sejla Goletic, Toni Eterovic, Adis Softic, Aida Kustura, Lana Salihefendic, Maja Ostojic, Maja Travar, Visnja Mrdjen, Nijaz Tihic, Sead Jazic, Sanjin Musa, Damir Marjanovic, Mirsada Hukic

**Affiliations:** Veterinary faculty of University of Sarajevo, Zmaja od Bosne 90, 71000 Sarajevo, Bosnia and Herzegovina; ALEA Genetic Center, Olovska 67, 71000 Sarajevo, Bosnia and Herzegovina; University Clinical Hospital of Mostar, Bijeli Brijeg b.b., 88000 Mostar, Bosnia and Herzegovina; University Clinical Centre of the Republic of Srpska, Dvanaest beba bb, 78000 Banja Luka, Bosnia and Herzegovina; University Clinical Center Tuzla, Ulica Profesora doktora Ibre Pašića, 75000 Tuzla, Bosnia and Herzegovina; General Hospital “Abdulah Nakas”, Kranjceviceva 12, 71000 Sarajevo, Bosnia and Herzegovina; Institute for Public Health of Federation of Bosnia and Herzegovina, Marsala Tita 9, 71000 Sarajevo, Bosnia and Herzegovina; Center for Applied Bioanthropology, Institute for Anthropological Researches, Gajeva ulica 32, 10 000 Zagreb, Croatia; Department of Genetics and Bioengineering, Faculty of Engineering and Natural Sciences, International Burch University, Francuske revolucije bb, 71 000 Sarajevo, Bosnia and Herzegovina; The Academy of Science and Arts of Bosnia and Herzegovina, Bistrik 7, 71000 Sarajevo, Bosnia and Herzegovina

## Abstract

Whole Genome Sequence of four samples from COVID-19 outbreaks was done in two laboratories in Bosnia and Herzegovina (Veterinary Faculty Sarajevo and Alea Genetic Center). All four BiH sequences cluster mainly with European ones (Italy, Austria, France, Sweden, Cyprus, England). The constructed phylogenetic tree indicates probable multiple independent introduction events. The success of future containment measures concernig new introductions will be highly challenging for country due to the significant proportion of BH population living abroad.

## Background and objectives

The first case of a new viral respiratory disease (COVID-19) caused by a newly discovered virus (SARS-CoV-2) was confirmed in Bosnia and Herzegovina (BH) on March 5, 2020. In the period until 17^th^ of June, a total of 3085 cases were laboratory confirmed, of which 168 with fatal outcome [1]. On 17^th^ of March 2020 BH declared national emergency followed by introduction of very restrictive measures on international movement, social distancing, mandatory use of personal protective equipment and lockdown for the population under 18 and above 65 years. Despite having 13 different health authorities and no state ministry of health due to complex governmental structure, it is generally agreed that BH authorities successfully implemented case detecting and tracing practices keeping epidemic scenario not to overwhelm health care capacities. Cease of restrictive measures started mid-May 2020 and, regardless of the current new cases numbers increase in the first half of June, authorities are keeping the plan to return to pre epidemic life conditions. BH scientific community has a significant experience in, and previously established capacity for, a wide specter of both human and animal virologic genetic analysis [2,3,4,5]. As a result, several scientific teams were prepared to be involved in a joint mission of genome sequencing of SARS-CoV-2 in BH. Objectives of this research were:

To share obtained sequences of the complete genome of SARS-CoV-2 strains from clinical samples of BiH patients diagnosed with COVID-19, and To contribute to the understanding of the interaction of molecular and classical epidemiology findings of COVID 19 in BH and the whole region and give recommendations for the improvement of prevention and future measures.

### Samples, RNA extraction and whole genome sequencing (WGS)

The samples used in this study were nasopharyngeal swabs in Viral Transport Medium from four patients, originating from Tuzla, Sarajevo, Livno and Banja Luka, and RT-PCR COVID-19 confirmed on 6, 8, 11, and 29 of April 2020, respectively (Figure 1.).

**Figure 1.**
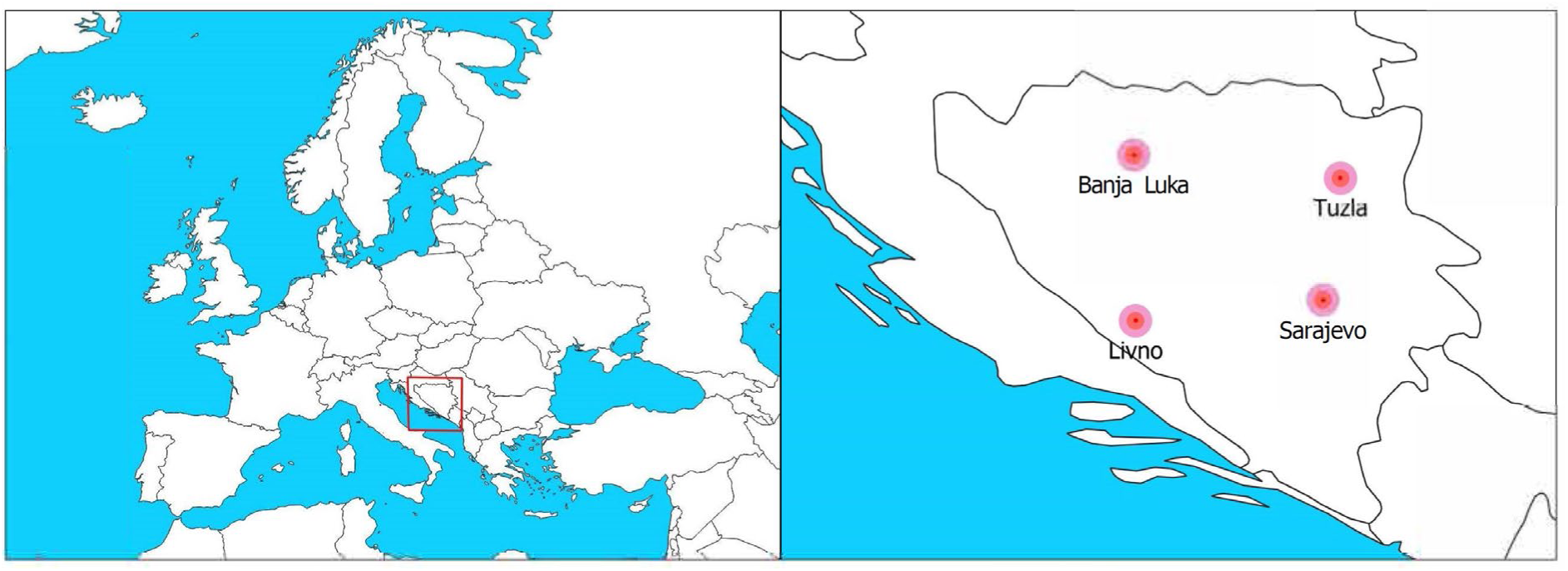
Sample locations in BH (Livno, Banja Luka, Tuzla, Sarajevo). Description: samples were collected from four geographically representative different regions in BH.

Samples from Livno and Banja Luka were processed at the laboratory of Veterinary Faculty Sarajevo (VFS) and samples from Tuzla and Sarajevo at the Alea Genetic Center, Sarajevo (AGC). A total of 140 μL of each sample was used for RNA extraction by QIAamp Viral RNA Mini Kit (Qiagen, Germany), and the product of extraction was later used for further analysis, while the rest of the samples were used for virus isolation on various cell lines at the VFS (data not shown here). The presence of SARS-CoV-2 in the samples was confirmed by real-time RT-PCR using HKU protocol [6], and the cycle threshold (Ct) values were 17,1; 21,3; 24,6 and 20,4 for Livno, Banja Luka, Tuzla and Sarajevo samples, respectively.

#### Livno and Banja Luka samples

WGS was performed according to the ARTIC amplicon sequencing protocol for MinION for nCoV-2019, which uses two primer pools to generate the sequence, as described elsewhere [7]. The sequencing was performed on a MinION sequencer, using an R9.4.1 flow cell on which the samples, as well as a negative control, were pooled. MinKNOW software and the MinIT device were used for high accuracy real-time base-calling during the run, which lasted 17 hours. Only the base-called fastq files with the Q score ≥ 7 were used for further analysis. The bioinformatic analysis was performed according to the nCoV-2019 novel coronavirus bioinformatics protocol [8]. The consensus sequence was mapped, for correction purposes, to the Wuhan reference genome (MN908947) using Minimap2, and polished in Racon.

#### Tuzla and Sarajevo samples

WGS was performed according to Ion AmpliSeq™ SARS-CoV-2 Research Panel Instructions for use on an Ion GeneStudio™ S5 Series System, as described previously [9], with certain modifications. Sequencing was performed on Ion GeneStudio™ S5 instrument using 520 chip. Raw data was analyzed using Torrent Suite Software 5.12.0 where the sequences were aligned to the Ion AmpliSeq SARS-CoV-2 reference genome. FASTA format of the obtained sequences was generated by the Iterative Refinement Meta-Assembler (IRMA) plugin. Sequences were additionally reviewed using BioEdit software.

All sequences were deposited in the Global Initiative on Sharing All Influenza Data (GISAID) https://www.gisaid.org/epiflu-applications/next-hcov-19-app/).

### Phylogenetic analysis

To perform a phylogenetic analysis (PA) of sequences from Livno (GISAID Accession ID: EPI_ISL_462753), Banja Luka (GISAID Accession ID: EPI_ISL_462990), Tuzla (GISAID Accession ID: EPI_ISL_463893) and Sarajevo (GISAID Accession ID: EPI_ISL_467300), a dataset of 120 whole genome sequences was obtained from GISAID (Supplementary material). These sequences were chosen based on their similarity with four BH sequences mentioned above. Namely, a set of 30 sequences from other countries, displaying the highest sequence identity with each BH sequence, were chosen. Sequence alignment and the construction of the phylogenetic tree were performed in MEGA X [10].

Two main GISAID lineages [11] were highlighted with differently coloured brackets and named accordingly: B.1.1. (GR) (yellow bracket) and B.1 (G) (blue bracket). Four BH sequences were marked with a different marker (red square, circles and a triangle), and bootstrap values were shown at the level of nodes. Livno sequence (GISAID Accession ID: EPI_ISL_462753) is marked differently from other three BH sequences because its GISAID subclade (B.1.1. (O)) is different than the subclade it has been sorted in this phylogenetic tree (B.1.1. (GR)).

Position 614 in Spike protein, used to characterize the G clade [12], was shown to be G in BH isolates, assigning all four BiH isolates to G clade together with European sequences (Italy, Austria, France, Sweden, Cyprus, England). The constructed phylogenetic tree in Figure 2 indicates probable multiple independent introduction events as reflected by clustering of each single BiH sequence in a separate cluster, highlighted with red (Livno, EPI_ISL_462753), green (Banja Luka, EPI_ISL_462990), blue (Sarajevo, EPI_ISL_467300) and purple (Tuzla, EPI_ISL_463893).

**Figure 2.**
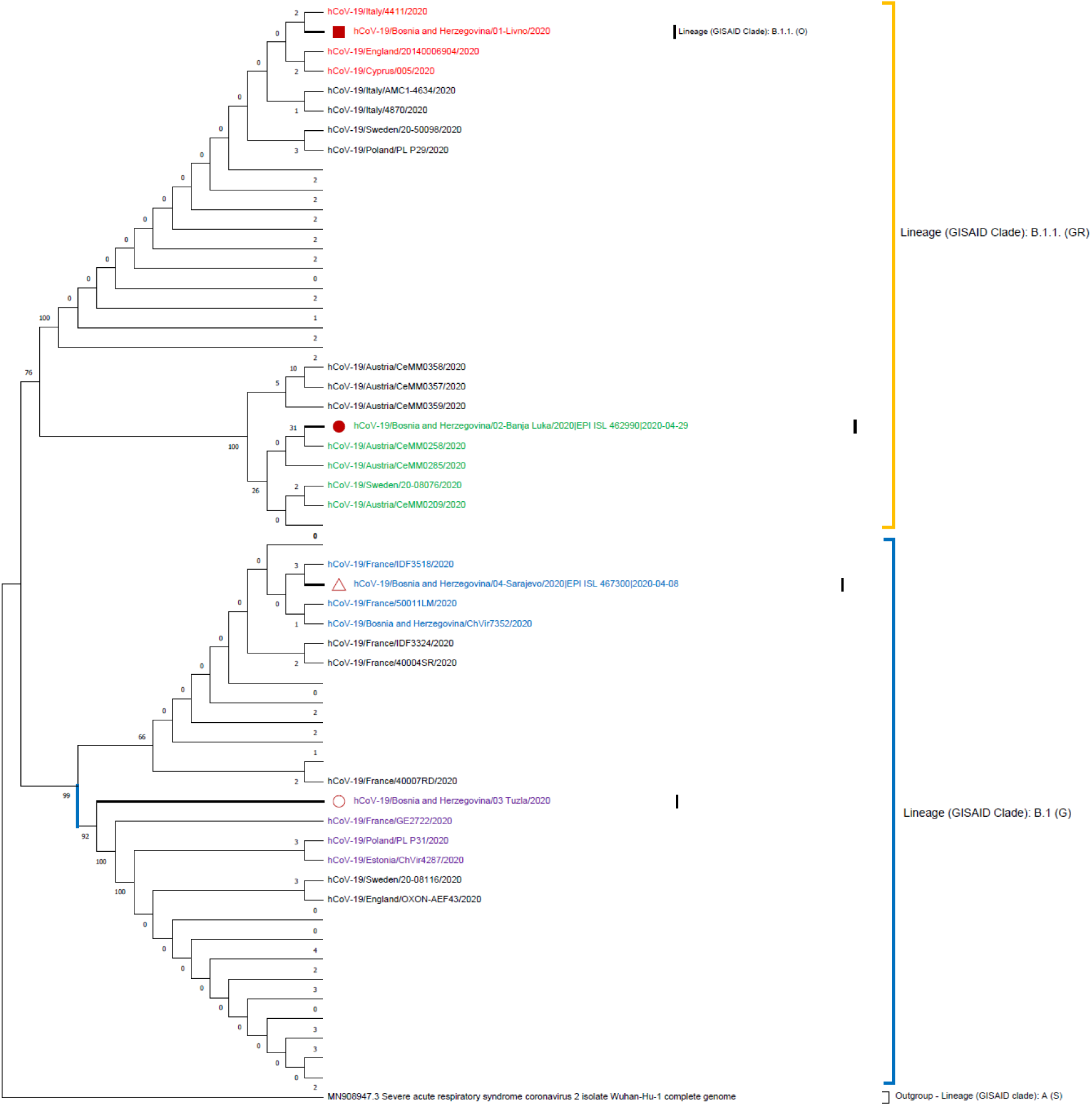
Maximum likelihood phylogenetic tree of four SARS-CoV-2 genome sequences retrieved in this study, as well as sequences from different countries (n = 120 genome sequences). Description: The phylogenetic tree was created using the Maximum Likelihood method, Hasegawa-Kishino-Yano substitution model as well as 1000 bootstrap replicates. Main clusters are highlighted in different colours. Certain branches have been collapsed and labels have been deleted for the ease of browsing.

## Discussion and conclusions

We sequenced the whole genome out of four samples from four different BH regions, originated from Tuzla, Livno, Sarajevo and Banjaluka municipalities. Even though all four BiH sequences cluster mainly with European ones, the constructed phylogenetic tree indicates multiple independent introductions of COVID-19 in BiH. Those findings correspond with other COVID-19 WGS studies [13,14] and confirm the importance of travel-associated disease introduction events for BiH as well. The success of future mitigation measures related to this type of disease introduction will be highly challenging for BiH due to a significant proportion of BiH population living abroad. WGS is a valuable approach for further investigation of the level of association of various modes of international movement (touristic, economic, religious, family visits etc) with transmission chains and associated risks.

Recently, significant increase in the number of new cases (more than 280 in just one week and both international and community transmitted) emphasized a need to establish a more efficient scientific and expert system for further work on the genetic analysis of the SARS CoV-2 virus, and to continue investigation of the whole genetic COVID-19 pattern as well. Integration of phylogenetic (molecular) and epidemiological approaches in the assessment of human, animal and environmental data will help with the identification of risk factors for disease spreading and optimize efficient and rational use of preventive measures. The importance of results gained from such approaches and studies will help in communication and scientific justification of social distancing and movement restriction perception in public.

### Data availability

All generated viral genome assemblies have been submitted to GSIAD. All submitters of data may be contacted directly via www.gisaid.org

## Supporting information

List of sequences obtained from GISAID and uncollapsed phylogenetic tree

## Acknowledgements

Part of this work was conducted in molecular-diagnostic and forensic laboratory of Veterinary faculty University of Sarajevo that is supported with equipment and training of staff by International Atomic Energy Agency (IAEA) under project “Strengthening State Infrastructure for Food and Animal Food Control and Protecting Animal Health” (BOH 5002). The authors wish to thank Dr Joshua Quick for his kind contribution of ARTIC primer set for whole genome sequencing, which greatly assisted in their research. The authors wish to thank Dr Goran Cerkez from Federal ministry of health for his continuing support of this research and Dr Mirza Ponjavic from University of Tuzla for technical support. We gratefully acknowledge the Authors, the Originating and Submitting laboratories for their sample and metadata shared.

## Ethical statement

Our study was approved by the ethics committee of Veterinary faculty of University of Sarajevo (#01-02-18-17/2020).

## Authors’ contribution

TG, RK, DM and NF conceptualised and designed the study. TG, RK, SG, TE, AK, AS, and LS designed and implemented lab protocols. TG, RK and SG developed and implemented sequencing data analysis approaches. MH, MO, MT, VM, NT, SJ and SM provided clinical samples and epidemiological data. TG, RK, SG, DM, SM and NF made an analysis of data and drafted discussion and conclusions of the study. All authors have commented on the draft and approved the final version.

## References

1. Anonymous. Ministry of Civil Affairs of Bosnia and Herzegovina. http://mcp.gov.ba/publication/read/epidemioloska-slika-covid-19?pageId=3 (accessed on 17 of June 2020)

2. Marjanovic D, Durmic-Pasic A, Bakal N, Haveric S, Kalamujic B, Kovacevic L, et al. DNA Identification of Skeletal Remains from the Second World War Mass Graves Uncovered in Slovenia. Croat Med J. 2007;48(4):513–9.

3. Davoren J, Vanek D, Konjhodzic R, Crews J, Huffine E, Parson TJ. Highly Effective DNA Extraction Method for Nuclear Short Tandem Repeat Testing of Skeletal Remains from Mass Graves. Croat Med J. 2007;48(4):478–85.

4. Marjanovic D, Hadzic Metjahic N, Cakar J, Dzehverovic M, Dogan S, Feric E, et al. Identification of human remains from the Second World War mass graves uncovered in Bosnia and Herzegovina. Croat Med J. 2015;56(3):257–62.

5. Goletic T, Kustura A, Gagic A, Savic A, Residbegovic E, Kavazovic A, et al. Molecular determinants of pathogenicity and host specificity of highly pathogenic H5N1 BiH isolates. Acta Med Salin. 2019;49(1):25–33.

6. Poon L, Chu D, Peiris M. HKU protocol. School of Public Health, The University of Hong Kong, Hong Kong. WHO [Internet]. 2020 [cited 2020 Jun 14]. Available from: https://www.who.int/docs/default-source/coronaviruse/peiris-protocol-16-1-20.pdf?sfvrsn=af1aac73_4

7. Quick, J. nCoV-2019 sequencing protocol v2. Protocols.io [Internet]. 2020 Apr [cited 2020 Jun 14]. Available from: https://dx.doi.org/10.17504/protocols.io.bdp7i5rn

8. Loman N, Rowe W, Rambaut A. nCoV-2019 novel coronavirus bioinformatics protocol. Artic Network [Internet]. 2020 Jan [cited 2020 Jun 14]. Available from: https://artic.network/ncov-2019/ncov2019-bioinformatics-sop.html

9. Ion AmpliSeq™ SARS-CoV-2 Research Panel Instructions for use on an Ion GeneStudio™ S5 Series System Quick Reference (Pub. No. MAN0019277). Ampliseq.com [Internet]. 2020 Apr [cited 2020 Jun 14]. Available from: https://assets.thermofisher.com/TFSAssets/LSG/manuals/MAN0019277_Ion_AmpliSeq_SARS-CoV-2_Research_Panel_GeneStudio_QR.pdf

10. Kumar S, Stecher G, Li M, Knyaz C, Tamura K. MEGA X: Molecular Evolutionary Genetics Analysis across computing platforms. Mol Biol Evol. 2018;35(6):1547–9. https://doi.org/10.1093/molbev/msy096 PMID: 29722887

11. Rambaut A, Holmes EC, Hill V, O’Toole Á, McCrone JT, Ruis C, et al. A dynamic nomenclature proposal for SARS-CoV-2 to assist genomic epidemiology. BioRxiv.org [Internet]. 2020 Apr [cited 2020 Jun 14]. Available from: https://doi.org/10.1101/2020.04.17.046086

12. Mondal M, Lawarde A, Somasundaram K.: Genomics of Indian SARS-CoV-2: Implications in genetic diversity, possible origin and spread of virus. MedRxiv.org [Internet]. 2020 Apr [cited 2020 Jun 14]. Available from: https://doi.org/10.1101/2020.04.25.20079475

13. Walker A, Houwaart T, Wienemann T, Vasconcelos Malte K, Strelow D, Senff Tin, et al. Genetic structure of SARS-CoV-2 reflects clonal superspreading and multiple independent introduction events, North-Rhine Westphalia, Germany, February and March 2020. EuroSurveillance 2020;25(22) https://doi.org/10.2807/1560-7917.ES.2020.25.22.2000746

14. Filipe A, Shepherd J, Williams T, Hughes J, Aranday Cortes E, Asamaphan P, et al. Genomic epidemiology of SARS-CoV-2 spread in Scotland highlights the role of European travel in COVID-19 emergence. medRxiv 2020.06.08.20124834; doi: https://doi.org/10.1101/2020.06.08.20124834

