## Supplementary material for "Phylogenetic pattern of SARS-CoV-2 from COVID-19 patients from Bosnia and Herzegovina: lessons learned to optimize future molecular and epidemiological approaches": List of sequences obtained from GISAID and uncollapsed phylogenetic tree

This supplementary material is hosted by *Eurosurveillance* as supporting information alongside the article [Phylogenetic pattern of SARS-CoV-2 from COVID-19 patients from Bosnia and Herzegovina: what have we learned to optimize future molecular and epidemiological findings], on behalf of the authors, who remain responsible for the accuracy and appropriateness of the content. The same standards for ethics, copyright, attributions and permissions as for the article apply. Supplements are not edited by *Eurosurveillance* and the journal is not responsible for the maintenance of any links or email addresses provided therein.

The authors gratefully acknowledge the Authors, the Originating and Submitting laboratories for their sample and metadata shared. The accession IDs, names, submission dates, sample location, submitting laboratories and the authors of the sequences used in phylogenetic analysis can be found on GISAID and are listed below.

- EPI_ISL_438045; hCoV-19/Austria/CeMM0206/2020; 2020-05-12; Europe / Austria / Vienna; Bergthaler laboratory, CeMM Research Center for Molecular Medicine of the Austrian Academy of Sciences; Alexandra Popa, Benedikt Agerer, Henrique Colaco, Lukas Endler, Jakob-Wendelin Genger, Alexander Lercher, Mark Smyth, Thomas Penz, Michael Schuster, Jan Laine, Martin Senekowitsch, Judith Aberle, Stephan Aberle, Elisabeth Puchhammer-Stoeckl, Manfred Nairz, Guenter Weiss, Wegene Borena, Dorothee von Laer, Christoph Bock, Andreas Bergthaler.
- EPI_ISL_438046; hCoV-19/Austria/CeMM0209/2020; 2020-05-12; Europe / Austria / Vienna; Bergthaler laboratory, CeMM Research Center for Molecular Medicine of the Austrian Academy of Sciences; Alexandra Popa, Benedikt Agerer, Henrique Colaco, Lukas Endler, Jakob-Wendelin Genger, Alexander Lercher, Mark Smyth, Thomas Penz, Michael Schuster, Jan Laine, Martin Senekowitsch, Judith Aberle, Stephan Aberle, Elisabeth Puchhammer-Stoeckl, Manfred Nairz, Guenter Weiss, Wegene Borena, Dorothee von Laer, Christoph Bock, Andreas Bergthaler.
- EPI_ISL_438087; hCoV-19/Austria/CeMM0258/2020; 2020-05-12; Europe / Austria / Vienna; Bergthaler laboratory, CeMM Research Center for Molecular Medicine of the Austrian Academy of Sciences; Alexandra Popa, Benedikt Agerer, Henrique Colaco, Lukas Endler, Jakob-Wendelin Genger, Alexander Lercher, Mark Smyth, Thomas Penz, Michael Schuster, Jan Laine, Martin Senekowitsch, Judith Aberle, Stephan Aberle, Elisabeth Puchhammer-Stoeckl, Manfred Nairz, Guenter Weiss, Wegene Borena, Dorothee von Laer, Christoph Bock, Andreas Bergthaler.
- EPI_ISL_438111; hCoV-19/Austria/CeMM0285/2020; 2020-05-12; Europe / Austria / Vienna; Bergthaler laboratory, CeMM Research Center for Molecular Medicine of the Austrian Academy of Sciences; Alexandra Popa, Benedikt Agerer, Henrique Colaco, Lukas Endler, Jakob-Wendelin Genger, Alexander Lercher, Mark Smyth, Thomas Penz, Michael Schuster, Jan Laine, Martin Senekowitsch, Judith Aberle, Stephan Aberle, Elisabeth Puchhammer-Stoeckl, Manfred Nairz, Guenter Weiss, Wegene Borena, Dorothee von Laer, Christoph Bock, Andreas Bergthaler.
- EPI_ISL_438120; hCoV-19/Austria/CeMM0357/2020; 2020-05-12; Europe / Austria / Vienna; Bergthaler laboratory, CeMM Research Center for Molecular Medicine of the Austrian Academy of Sciences; Alexandra Popa, Benedikt Agerer, Henrique Colaco, Lukas Endler, Jakob-Wendelin Genger, Alexander Lercher, Mark Smyth, Thomas Penz, Michael Schuster, Jan Laine, Martin Senekowitsch, Judith Aberle, Stephan Aberle, Elisabeth Puchhammer-Stoeckl, Manfred Nairz, Guenter Weiss, Wegene Borena, Dorothee von Laer, Christoph Bock, Andreas Bergthaler.
- EPI_ISL_438121; hCoV-19/Austria/CeMM0358/2020; 2020-05-12; Europe / Austria / Vienna; Bergthaler laboratory, CeMM Research Center for Molecular Medicine of the Austrian Academy of Sciences; Alexandra Popa, Benedikt Agerer, Henrique Colaco, Lukas Endler, Jakob-Wendelin Genger, Alexander Lercher, Mark Smyth, Thomas Penz, Michael Schuster, Jan Laine, Martin Senekowitsch, Judith Aberle, Stephan Aberle, Elisabeth Puchhammer-Stoeckl, Manfred Nairz, Guenter Weiss, Wegene Borena, Dorothee von Laer, Christoph Bock, Andreas Bergthaler.
- EPI_ISL_438122; hCoV-19/Austria/CeMM0359/2020; 2020-05-12; Europe / Austria / Vienna; Bergthaler laboratory, CeMM Research Center for Molecular Medicine of the Austrian Academy of Sciences; Alexandra Popa, Benedikt Agerer, Henrique Colaco, Lukas Endler, Jakob-Wendelin Genger, Alexander Lercher, Mark Smyth, Thomas Penz, Michael Schuster, Jan Laine, Martin Senekowitsch, Judith Aberle, Stephan Aberle, Elisabeth Puchhammer-Stoeckl, Manfred Nairz, Guenter Weiss, Wegene Borena, Dorothee von Laer, Christoph Bock, Andreas Bergthaler.

- EPI_ISL_462990; hCoV-19/Bosnia and Herzegovina/02-Banja Luka/2020; 2020-04-29; Europe / Bosnia and Herzegovina / Čelinac; University of Sarajevo, Veterinary Faculty; Teufik, G., Šejla, G., Toni, E., Maja, T., Mirsada, H., Aida, K., Alma, Š.A.
- EPI_ISL_465868; hCoV-19/England/20140002404/2020; 2020-03-27; Europe / United Kingdom / England; Respiratory Virus Unit, Microbiology Services Colindale, Public Health England; PHE Covid Sequencing Team.
- EPI_ISL_454438; hCoV-19/Sweden/20-08043/2020; 2020-05-28; Europe / Sweden / Halland; The Public Health Agency of Sweden; Anna-Malin Linde, Maria Lind Karlberg, Mattias Haukland, Reza Advani, Olov Svartstrom, Oskar Karlsson Lindsjo, Petra Edquist, Shamam Muradrasoli, Anna Risberg, Karin Tegmark-Wisell.
- EPI_ISL_454447; hCoV-19/Sweden/20-08060/2020; 2020-05-28; Europe / Sweden / Gavleborg; The Public Health Agency of Sweden; Anna-Malin Linde, Maria Lind Karlberg, Mattias Haukland, Reza Advani, Olov Svartstrom, Oskar Karlsson Lindsjo, Petra Edquist, Shamam Muradrasoli, Anna Risberg, Karin Tegmark-Wisell.
- EPI_ISL_454451; hCoV-19/Sweden/20-08076/2020; 2020-05-28; Europe / Sweden / Stockholm; The Public Health Agency of Sweden; Anna-Malin Linde, Maria Lind Karlberg, Mattias Haukland, Reza Advani, Olov Svartstrom, Oskar Karlsson Lindsjo, Petra Edquist, Shamam Muradrasoli, Anna Risberg, Karin Tegmark-Wisell.
- EPI_ISL_454452; hCoV-19/Sweden/20-08077/2020; 2020-05-28; Europe / Sweden / Stockholm; The Public Health Agency of Sweden; Anna-Malin Linde, Maria Lind Karlberg, Mattias Haukland, Reza Advani, Olov Svartstrom, Oskar Karlsson Lindsjo, Petra Edquist, Shamam Muradrasoli, Anna Risberg, Karin Tegmark-Wisell.
- EPI_ISL_454453; hCoV-19/Sweden/20-08078/2020; 2020-05-28; Europe / Sweden / Stockholm; The Public Health Agency of Sweden; Anna-Malin Linde, Maria Lind Karlberg, Mattias Haukland, Reza Advani, Olov Svartstrom, Oskar Karlsson Lindsjo, Petra Edquist, Shamam Muradrasoli, Anna Risberg, Karin Tegmark-Wisell.
- EPI_ISL_454461; hCoV-19/Sweden/20-08088/2020; 2020-05-28; Europe / Sweden / Stockholm; The Public Health Agency of Sweden; Anna-Malin Linde, Maria Lind Karlberg, Mattias Haukland, Reza Advani, Olov Svartstrom, Oskar Karlsson Lindsjo, Petra Edquist, Shamam Muradrasoli, Anna Risberg, Karin Tegmark-Wisell.
- EPI_ISL_454870; hCoV-19/Sweden/20-50055/2020; 2020-05-29; Europe / Sweden / Stockholm; The Public Health Agency of Sweden; Anna-Malin Linde, Maria Lind Karlberg, Mattias Haukland, Reza Advani, Olov Svartstrom, Oskar Karlsson Lindsjo, Petra Edquist, Shamam Muradrasoli, Anna Risberg, Karin Tegmark-Wisell.
- EPI_ISL_455851; hCoV-19/Sweden/20-50095/2020; 2020-06-01; Europe / Sweden / Stockholm; The Public Health Agency of Sweden; Anna-Malin Linde, Maria Lind Karlberg, Mattias Haukland, Reza Advani, Olov Svartstrom, Oskar Karlsson Lindsjo, Petra Edquist, Shamam Muradrasoli, Anna Risberg, Karin Tegmark-Wisell.
- EPI_ISL_454881; hCoV-19/Sweden/20-50107/2020; 2020-05-29; Europe / Sweden / Stockholm; The Public Health Agency of Sweden; Anna-Malin Linde, Maria Lind Karlberg, Mattias Haukland, Reza Advani, Olov Svartstrom, Oskar Karlsson Lindsjo, Petra Edquist, Shamam Muradrasoli, Anna Risberg, Karin Tegmark-Wisell.
- EPI_ISL_454883; hCoV-19/Sweden/20-50139/2020; 2020-05-29; Europe / Sweden / Stockholm; The Public Health Agency of Sweden; Anna-Malin Linde, Maria Lind Karlberg, Mattias Haukland, Reza Advani, Olov Svartstrom, Oskar Karlsson Lindsjo, Petra Edquist, Shamam Muradrasoli, Anna Risberg, Karin Tegmark-Wisell.
- EPI_ISL_455860; hCoV-19/Sweden/20-50150/2020; 2020-06-01; Europe / Sweden / Stockholm; The Public Health Agency of Sweden; Anna-Malin Linde, Maria Lind Karlberg, Mattias Haukland, Reza Advani, Olov Svartstrom, Oskar Karlsson Lindsjo, Petra Edquist, Shamam Muradrasoli, Anna Risberg, Karin Tegmark-Wisell.
- EPI_ISL_455871; hCoV-19/Sweden/20-50212/2020; 2020-06-01; Europe / Sweden / Stockholm; The Public Health Agency of Sweden; Anna-Malin Linde, Maria Lind Karlberg, Mattias Haukland, Reza Advani, Olov Svartstrom, Oskar Karlsson Lindsjo, Petra Edquist, Shamam Muradrasoli, Anna Risberg, Karin Tegmark-Wisell.
- EPI_ISL_455874; hCoV-19/Sweden/20-50217/2020; 2020-06-01; Europe / Sweden / Stockholm; The Public Health Agency of Sweden; Anna-Malin Linde, Maria Lind Karlberg,

Mattias Haukland, Reza Advani, Olov Svartstrom, Oskar Karlsson Lindsjo, Petra Edquist, Shamam Muradrasoli, Anna Risberg, Karin Tegmark-Wisell.

- EPI_ISL_455875; hCoV-19/Sweden/20-50218/2020; 2020-06-01; Europe / Sweden / Stockholm; The Public Health Agency of Sweden; Anna-Malin Linde, Maria Lind Karlberg, Mattias Haukland, Reza Advani, Olov Svartstrom, Oskar Karlsson Lindsjo, Petra Edquist, Shamam Muradrasoli, Anna Risberg, Karin Tegmark-Wisell.
- EPI_ISL_455879; hCoV-19/Sweden/20-50229/2020; 2020-06-01; Europe / Sweden / Stockholm; The Public Health Agency of Sweden; Anna-Malin Linde, Maria Lind Karlberg, Mattias Haukland, Reza Advani, Olov Svartstrom, Oskar Karlsson Lindsjo, Petra Edquist, Shamam Muradrasoli, Anna Risberg, Karin Tegmark-Wisell.
- EPI_ISL_455892; hCoV-19/Sweden/20-50254/2020; 2020-06-01; Europe / Sweden / Stockholm; The Public Health Agency of Sweden; Anna-Malin Linde, Maria Lind Karlberg, Mattias Haukland, Reza Advani, Olov Svartstrom, Oskar Karlsson Lindsjo, Petra Edquist, Shamam Muradrasoli, Anna Risberg, Karin Tegmark-Wisell.
- EPI_ISL_455893; hCoV-19/Sweden/20-50255/2020; 2020-06-01; Europe / Sweden / Stockholm; The Public Health Agency of Sweden; Anna-Malin Linde, Maria Lind Karlberg, Mattias Haukland, Reza Advani, Olov Svartstrom, Oskar Karlsson Lindsjo, Petra Edquist, Shamam Muradrasoli, Anna Risberg, Karin Tegmark-Wisell.
- EPI_ISL_455894; hCoV-19/Sweden/20-50256/2020; 2020-06-01; Europe / Sweden / Stockholm; The Public Health Agency of Sweden; Anna-Malin Linde, Maria Lind Karlberg, Mattias Haukland, Reza Advani, Olov Svartstrom, Oskar Karlsson Lindsjo, Petra Edquist, Shamam Muradrasoli, Anna Risberg, Karin Tegmark-Wisell.
- EPI_ISL_455897; hCoV-19/Sweden/20-50259/2020; 2020-06-01; Europe / Sweden / Stockholm; The Public Health Agency of Sweden; Anna-Malin Linde, Maria Lind Karlberg, Mattias Haukland, Reza Advani, Olov Svartstrom, Oskar Karlsson Lindsjo, Petra Edquist, Shamam Muradrasoli, Anna Risberg, Karin Tegmark-Wisell.
- EPI_ISL_455904; hCoV-19/Sweden/20-50270/2020; 2020-06-01; Europe / Sweden / Stockholm; The Public Health Agency of Sweden; Anna-Malin Linde, Maria Lind Karlberg, Mattias Haukland, Reza Advani, Olov Svartstrom, Oskar Karlsson Lindsjo, Petra Edquist, Shamam Muradrasoli, Anna Risberg, Karin Tegmark-Wisell.
- EPI_ISL_450197; hCoV-19/Thailand/Nonthaburi_2520/2020; 2020-05-20; Asia / Thailand / Nonthaburi; National Institute of Health. Department of medical Sciences, Ministry of Public Health, Thailand; Pilailuk,Okada; Siripaporn,Phuygun; Thanutsapa,Thanadachakul; Sittiporn,Parnmen;Warawan,Wongboot; Sunthareeya,Waicharoen; Malinee,Chittaganpitch.
- EPI_ISL_462753; hCoV-19/Bosnia and Herzegovina/01-Livno/2020; 2020-06-08; Europe / Bosnia and Herzegovina / Livno; University of Sarajevo Veterinary Faculty; Goletic, T., Softic, A., Goletic, S., Ostojic, M., Hukic, M., Eterovic, T., Seho-Alic, A.
- EPI_ISL_463741; hCoV-19/Cyprus/001/2020; 2020-06-10; Europe / Cyprus; Department of Molecular Virology, Cyprus Institute of Neurology and Genetics; Jan Richter, George Krashias, Christina Tryfonos, Stavros Bashiardes, Dana Koptides, Christina Christodoulou.
- EPI_ISL_463745; hCoV-19/Cyprus/005/2020; 2020-06-10; Europe / Cyprus; Department of Molecular Virology, Cyprus Institute of Neurology and Genetics; Jan Richter, George Krashias, Christina Tryfonos, Stavros Bashiardes, Dana Koptides, Christina Christodoulou.
- EPI_ISL_463746; hCoV-19/Cyprus/006/2020; 2020-06-10; Europe / Cyprus; Department of Molecular Virology, Cyprus Institute of Neurology and Genetics; Jan Richter, George Krashias, Christina Tryfonos, Stavros Bashiardes, Dana Koptides, Christina Christodoulou.
- EPI_ISL_464923; hCoV-19/England/20118050804/2020; 2020-06-11; Europe / United Kingdom / England; Respiratory Virus Unit, Microbiology Services Colindale, Public Health England; PHE Covid Sequencing Team.
- EPI_ISL_465875; hCoV-19/England/20140006904/2020; 2020-06-11; Europe / United Kingdom / England; Respiratory Virus Unit, Microbiology Services Colindale, Public Health England; PHE Covid Sequencing Team.
- EPI_ISL_452219; hCoV-19/Germany/FrankfurtFFM3/2020; 2020-05-27; Europe / Germany / Frankfurt; Widera/Toptan; Tuna Toptan1, Sebastian Hoehl1, Sandra Westhaus1, Denisa Bojkova1, Annemarie Berger1, Björn Rotter2, Klaus Hoffmeier2, Jindrich Cinatl1, Sandra Ciesek1, and Marek Widera.

- EPI_ISL_457728; hCoV-19/Italy/2389/2020; 2020-06-02; Europe / Italy / Molise; Army Medical and Veterinary Research Center; Paola Stefanelli, Alessandra Lo Presti, Stefano Fiore, Antonella Marchi, Eleonora Benedetti, Concetta Fabiani Silvia Fillo, Giovanni Faggioni, Riccardo De Sanctis, Antonella Fortunato, Anna Anselmo, Francesco Giordani, Vanessa Vera Fain, Nino D'Amore, Florigio Lista.
- EPI_ISL_457721; hCoV-19/Italy/4411/2020; 2020-06-02; Europe / Italy / Sicily; Army Medical and Veterinary Research Center; Paola Stefanelli, Alessandra Lo Presti, Stefano Fiore, Antonella Marchi, Eleonora Benedetti, Concetta Fabiani Silvia Fillo, Giovanni Faggioni, Riccardo De Sanctis, Antonella Fortunato, Anna Anselmo, Francesco Giordani, Vanessa Vera Fain, Nino D'Amore, Florigio Lista.
- EPI_ISL_457732; hCoV-19/Italy/4870/2020; 2020-06-02; Europe / Italy / Valle D'Aosta; Army Medical and Veterinary Research Center; Paola Stefanelli, Alessandra Lo Presti, Stefano Fiore, Antonella Marchi, Eleonora Benedetti, Concetta Fabiani Silvia Fillo, Giovanni Faggioni, Riccardo De Sanctis, Antonella Fortunato, Anna Anselmo, Francesco Giordani, Vanessa Vera Fain, Nino D'Amore, Florigio Lista.
- EPI_ISL_457736; hCoV-19/Italy/5062/2020; 2020-06-02; Europe / Italy / Basilicata; Army Medical and Veterinary Research Center; Paola Stefanelli, Alessandra Lo Presti, Stefano Fiore, Antonella Marchi, Eleonora Benedetti, Concetta Fabiani Silvia Fillo, Giovanni Faggioni, Riccardo De Sanctis, Antonella Fortunato, Anna Anselmo, Francesco Giordani, Vanessa Vera Fain, Nino D'Amore, Florigio Lista.
- EPI_ISL_457825; hCoV-19/Italy/AMC1-4634/2020; 2020-06-02; Europe / Italy / Sardinia; Army Medical and Veterinary Research Center; Silvia Fillo, Giovanni Faggioni, Riccardo De Sanctis, Antonella Fortunato, Anna Anselmo, Francesco Giordani, Vanessa Vera Fain, Nino D'Amore, Florigio Lista.
- EPI_ISL_455442; hCoV-19/Poland/PL_P29/2020; 2020-05-29; Europe / Poland / Mazowieckie / Grodzisk Mazowiecki; 1. ViroGenetics - BSL3 Laboratory of Virology, Małopolska Centre of Biotechnology, Jagiellonian University; 2. II Department of Internal Medicine, Faculty of Medicine, Jagiellonian University Medical College; 3. Narodowy Instytut Zdrowia Publicznego – Państwowy Zakład Higieny (NIZP-PZH); Katarzyna Pancer, Marek Sanak, Aleksandra A. Zasada, Magdalena Rzeczkowska, Tomasz Wołkowicz, Katarzyna Zacharczuk, Agnieszka Kołakowska-Kulesza, Katarzyna Owczarek, Aleksandra Milewska, Natalia Wolaniuk, Ewelina Hallman-Szelińska, Paweł P Łabaj, Wojciech Branicki, Krzysztof Pyrć.
- EPI_ISL_454473; hCoV-19/Sweden/20-08102/2020; 2020-05-28; Europe / Sweden / Stockholm; The Public Health Agency of Sweden; Anna-Malin Linde, Maria Lind Karlberg, Mattias Haukland, Reza Advani, Olov Svartstrom, Oskar Karlsson Lindsjo, Petra Edquist, Shamam Muradrasoli, Anna Risberg, Karin Tegmark-Wisell.
- EPI_ISL_454474; hCoV-19/Sweden/20-08103/2020; 2020-05-28; Europe / Sweden / Stockholm; The Public Health Agency of Sweden; Anna-Malin Linde, Maria Lind Karlberg, Mattias Haukland, Reza Advani, Olov Svartstrom, Oskar Karlsson Lindsjo, Petra Edquist, Shamam Muradrasoli, Anna Risberg, Karin Tegmark-Wisell.
- EPI_ISL_454486; hCoV-19/Sweden/20-08115/2020; 2020-05-28; Europe / Sweden / Stockholm; The Public Health Agency of Sweden; Anna-Malin Linde, Maria Lind Karlberg, Mattias Haukland, Reza Advani, Olov Svartstrom, Oskar Karlsson Lindsjo, Petra Edquist, Shamam Muradrasoli, Anna Risberg, Karin Tegmark-Wisell.
- EPI_ISL_454868; hCoV-19/Sweden/20-50052/2020; 2020-05-29; Europe / Sweden / Stockholm; The Public Health Agency of Sweden; Anna-Malin Linde, Maria Lind Karlberg, Mattias Haukland, Reza Advani, Olov Svartstrom, Oskar Karlsson Lindsjo, Petra Edquist, Shamam Muradrasoli, Anna Risberg, Karin Tegmark-Wisell.
- EPI_ISL_454869; hCoV-19/Sweden/20-50053/2020; 2020-05-29; Europe / Sweden / Stockholm; The Public Health Agency of Sweden; Anna-Malin Linde, Maria Lind Karlberg, Mattias Haukland, Reza Advani, Olov Svartstrom, Oskar Karlsson Lindsjo, Petra Edquist, Shamam Muradrasoli, Anna Risberg, Karin Tegmark-Wisell.
- EPI_ISL_454871; hCoV-19/Sweden/20-50088/2020; 2020-05-29; Europe / Sweden / Stockholm; The Public Health Agency of Sweden; Anna-Malin Linde, Maria Lind Karlberg, Mattias Haukland, Reza Advani, Olov Svartstrom, Oskar Karlsson Lindsjo, Petra Edquist, Shamam Muradrasoli, Anna Risberg, Karin Tegmark-Wisell.

- EPI_ISL_455850; hCoV-19/Sweden/20-50094/2020; 2020-06-01; Europe / Sweden / Stockholm; The Public Health Agency of Sweden; Anna-Malin Linde, Maria Lind Karlberg, Mattias Haukland, Reza Advani, Olov Svartstrom, Oskar Karlsson Lindsjo, Petra Edquist, Shamam Muradrasoli, Anna Risberg, Karin Tegmark-Wisell.
- EPI_ISL_454877; hCoV-19/Sweden/20-50098/2020; 2020-05-29; Europe / Sweden / Stockholm; The Public Health Agency of Sweden; Anna-Malin Linde, Maria Lind Karlberg, Mattias Haukland, Reza Advani, Olov Svartstrom, Oskar Karlsson Lindsjo, Petra Edquist, Shamam Muradrasoli, Anna Risberg, Karin Tegmark-Wisell.
- EPI_ISL_454879; hCoV-19/Sweden/20-50101/2020; 2020-05-29; Europe / Sweden / Stockholm; The Public Health Agency of Sweden; Anna-Malin Linde, Maria Lind Karlberg, Mattias Haukland, Reza Advani, Olov Svartstrom, Oskar Karlsson Lindsjo, Petra Edquist, Shamam Muradrasoli, Anna Risberg, Karin Tegmark-Wisell.
- EPI_ISL_455856; hCoV-19/Sweden/20-50140/2020; 2020-06-01; Europe / Sweden / Stockholm; The Public Health Agency of Sweden; Anna-Malin Linde, Maria Lind Karlberg, Mattias Haukland, Reza Advani, Olov Svartstrom, Oskar Karlsson Lindsjo, Petra Edquist, Shamam Muradrasoli, Anna Risberg, Karin Tegmark-Wisell.
- EPI_ISL_454891; hCoV-19/Sweden/20-50222/2020; 2020-05-29; Europe / Sweden / Stockholm; The Public Health Agency of Sweden; Anna-Malin Linde, Maria Lind Karlberg, Mattias Haukland, Reza Advani, Olov Svartstrom, Oskar Karlsson Lindsjo, Petra Edquist, Shamam Muradrasoli, Anna Risberg, Karin Tegmark-Wisell.
- EPI_ISL_455887; hCoV-19/Sweden/20-50247/2020; 2020-06-01; Europe / Sweden / Stockholm; The Public Health Agency of Sweden; Anna-Malin Linde, Maria Lind Karlberg, Mattias Haukland, Reza Advani, Olov Svartstrom, Oskar Karlsson Lindsjo, Petra Edquist, Shamam Muradrasoli, Anna Risberg, Karin Tegmark-Wisell.
- EPI_ISL_455891; hCoV-19/Sweden/20-50253/2020; 2020-06-01; Europe / Sweden / Stockholm; The Public Health Agency of Sweden; Anna-Malin Linde, Maria Lind Karlberg, Mattias Haukland, Reza Advani, Olov Svartstrom, Oskar Karlsson Lindsjo, Petra Edquist, Shamam Muradrasoli, Anna Risberg, Karin Tegmark-Wisell.
- EPI_ISL_455895; hCoV-19/Sweden/20-50257/2020; 2020-06-01; Europe / Sweden / Stockholm; The Public Health Agency of Sweden; Anna-Malin Linde, Maria Lind Karlberg, Mattias Haukland, Reza Advani, Olov Svartstrom, Oskar Karlsson Lindsjo, Petra Edquist, Shamam Muradrasoli, Anna Risberg, Karin Tegmark-Wisell.
- EPI_ISL_452108; hCoV-19/USA/MN-CDC-0103/2020; 2020-05-26; North America / USA / Minnesota; Pathogen Discovery, Respiratory Viruses Branch, Division of Viral Diseases, Centers for Disease Control and Prevention; Yan Li, Anna Montmayeur, Ying Tao, Krista Queen, Jing Zhang, Anna Uehara, Clinton R. Paden, Rachel Marine, Mary S. Keckler, Alison S. Laufer Halpin, Haibin Wang, Christopher A. Elkins, Zachary Weiner, Suxiang Tong.
- EPI_ISL_456031; hCoV-19/USA/NY-NYUMC795/2020; 2020-06-01; North America / USA / New York / Brooklyn; Departments of Pathology and Medicine, New York University School of Medicine; Maria Aguero-Rosenfeld, Brendan Belovarac, Margaret Black, Ludovic Boytard, John Cadley, Paolo Cotzia, John Chen, Dacia Dimartino, Xiaojun Feng, Tatyana Gindin, Emily Guzman, Adriana Heguy, Megan Hogan, Emily Huang, George Jour, Alireza Khodadadi-Jamayran, Lawrence H. Lin, Raven Luther, Andrew Lytle, Christian Marier, Matthew T. Maurano, Mark J. Mulligan, Peter Meyn, Raquel Ordonez Ciriza, Iman Osman, Jared Pinnell, Vanessa Raabe, Sitharam Ramaswami, Amy Rapkiewicz, Andre M. Ribeiro-dos-Santos, Marie Samanovic-Golden, Antonio Serrano, Guomiao Shen, Matija Snuderl, Theodore Vougiouklakis, Nick Vulpescu, Gael Westby, Paul Zappile, Yutong Zhang.
- EPI_ISL_452121; hCoV-19/USA/VI-CDC-3661/2020; 2020-05-26; North America / USA / Virgin Islands; Pathogen Discovery, Respiratory Viruses Branch, Division of Viral Diseases, Centers for Disease Control and Prevention; Jing Zhang, Anna Montmayeur, Yan Li, Ying Tao, Krista Queen, Anna Uehara, Clinton R. Paden, Rachel Marine, Mary S. Keckler, Alison S. Laufer Halpin, Haibin Wang, Christopher A. Elkins, Zachary Weiner, Suxiang Tong.
- EPI_ISL_467300; hCoV-19/Bosnia and Herzegovina/04-Sarajevo/2020; 2020-06-12; Europe / Bosnia and Herzegovina / Sarajevo; Alea Genetic Center; Rijad Konjhodzic; Lana Salihefendic; Teufik Goletic; Sead Jazic; Dino Pecar; Nihad Fejzic; Damir Marjanovic; Enis Kandic.
- EPI_ISL_462451; hCoV-19/Bosnia and Herzegovina/ChVir7343/2020; 2020-06-08; Europe / Bosnia and Herzegovina / Bihac; Charite Universitatsmedizin Berlin, Institute of Virology; Victor

M Corman, Jorn Beheim-Schwarzbach, Barbara Muehlemann, Talitha Veith, Julia Schneider, Terry Jones, Amela Dedeic-Ljubovic, Irma Salimovic-Besic, Suzana Arapcic, Almedina Hadzihasanovic-Moro, Selma Mutevelic, Christian Drosten.

- EPI_ISL_462458; hCoV-19/Bosnia and Herzegovina/ChVir7352/2020; 2020-06-08; Europe / Bosnia and Herzegovina / Bihac; Charite Universitatsmedizin Berlin, Institute of Virology; Victor M Corman, Jorn Beheim-Schwarzbach, Barbara Muehlemann, Talitha Veith, Julia Schneider, Terry Jones, Amela Dedeic-Ljubovic, Irma Salimovic-Besic, Suzana Arapcic, Almedina Hadzihasanovic-Moro, Selma Mutevelic, Christian Drosten.
- EPI_ISL_450304; hCoV-19/Canada/Qc-L00240652/2020; 2020-05-20; North America / Canada / Quebec; Laboratoire de santé publique du Québec; Sandrine Moreira, Ioannis Ragoussis, Guillaume Bourque, Jesse Shapiro, Mark Lathrop and Michel Roger on behalf of the CoVSeQ research group (http://covseq.ca/researchgroup).
- EPI_ISL_445282; hCoV-19/Chile/Punta_Arenas_4/2020; 2020-05-15; South America / Chile / Punta Arenas; Instituto de Salud Publica de Chile; Andrés E Castillo, Bárbara Parra,Paz Tapia, Jaime Lagos, Loredana Arata, Alejandra Acevedo, Winston Andrade, Gabriel Leal, Carolina Tambley, Patricia Bustos, Rodrigo Fasce, Jorge Fernandez.
- EPI_ISL_447721; hCoV-19/France/10002PM/2020; 2020-05-14; Europe / France; Genomic platform; De Prost,N., Fourati,S., Lamoureux,C., Schmitz,D., Deveaux,I., Picard,O., Lepeule,R., Surgers,L., Mekontso-Dessap,A., Woerther,P.-L., Canoui-Poitrine,F., Pawlotsky,J.-M., Clinical Study Group,C., Rodrigue,C., Gricourt,G., N'debi,M., Demontant,V., Trawinski,E.
- EPI_ISL_447719; hCoV-19/France/10003SN/2020; 2020-05-14; Europe / France; Genomic platform; De Prost,N., Fourati,S., Lamoureux,C., Schmitz,D., Deveaux,I., Picard,O., Lepeule,R., Surgers,L., Mekontso-Dessap,A., Woerther,P.-L., Canoui-Poitrine,F., Pawlotsky,J.-M., Clinical Study Group,C., Rodrigue,C., Gricourt,G., N'debi,M., Demontant,V., Trawinski,E.
- EPI_ISL_447727; hCoV-19/France/10006HC/2020; 2020-05-14; Europe / France; Genomic platform; De Prost,N., Fourati,S., Lamoureux,C., Schmitz,D., Deveaux,I., Picard,O., Lepeule,R., Surgers,L., Mekontso-Dessap,A., Woerther,P.-L., Canoui-Poitrine,F., Pawlotsky,J.-M., Clinical Study Group,C., Rodrigue,C., Gricourt,G., N'debi,M., Demontant,V., Trawinski,E.
- EPI_ISL_447697; hCoV-19/France/10008DM/2020; 2020-05-14; Europe / France; Genomic platform; De Prost,N., Fourati,S., Lamoureux,C., Schmitz,D., Deveaux,I., Picard,O., Lepeule,R., Surgers,L., Mekontso-Dessap,A., Woerther,P.-L., Canoui-Poitrine,F., Pawlotsky,J.-M., Clinical Study Group,C., Rodrigue,C., Gricourt,G., N'debi,M., Demontant,V., Trawinski,E.
- EPI_ISL_447709; hCoV-19/France/10012BM/2020; 2020-05-14; Europe / France; Genomic platform; De Prost,N., Fourati,S., Lamoureux,C., Schmitz,D., Deveaux,I., Picard,O., Lepeule,R., Surgers,L., Mekontso-Dessap,A., Woerther,P.-L., Canoui-Poitrine,F., Pawlotsky,J.-M., Clinical Study Group,C., Rodrigue,C., Gricourt,G., N'debi,M., Demontant,V., Trawinski,E.
- EPI_ISL_447702; hCoV-19/France/10019LM/2020; 2020-05-14; Europe / France; Genomic platform; De Prost,N., Fourati,S., Lamoureux,C., Schmitz,D., Deveaux,I., Picard,O., Lepeule,R., Surgers,L., Mekontso-Dessap,A., Woerther,P.-L., Canoui-Poitrine,F., Pawlotsky,J.-M., Clinical Study Group,C., Rodrigue,C., Gricourt,G., N'debi,M., Demontant,V., Trawinski,E.
- EPI_ISL_447672; hCoV-19/France/10032KL/2020; 2020-05-14; Europe / France; Genomic platform; De Prost,N., Fourati,S., Lamoureux,C., Schmitz,D., Deveaux,I., Picard,O., Lepeule,R., Surgers,L., Mekontso-Dessap,A., Woerther,P.-L., Canoui-Poitrine,F., Pawlotsky,J.-M., Clinical Study Group,C., Rodrigue,C., Gricourt,G., N'debi,M., Demontant,V., Trawinski,E.
- EPI_ISL_447664; hCoV-19/France/10040LJ/2020; 2020-05-14; Europe / France; Genomic platform; De Prost,N., Fourati,S., Lamoureux,C., Schmitz,D., Deveaux,I., Picard,O., Lepeule,R., Surgers,L., Mekontso-Dessap,A., Woerther,P.-L., Canoui-Poitrine,F., Pawlotsky,J.-M., Clinical Study Group,C., Rodrigue,C., Gricourt,G., N'debi,M., Demontant,V., Trawinski,E.
- EPI_ISL_447687; hCoV-19/France/10078MA/2020; 2020-05-14; Europe / France; Genomic platform; De Prost,N., Fourati,S., Lamoureux,C., Schmitz,D., Deveaux,I., Picard,O., Lepeule,R., Surgers,L., Mekontso-Dessap,A., Woerther,P.-L., Canoui-Poitrine,F., Pawlotsky,J.-M., Clinical Study Group,C., Rodrigue,C., Gricourt,G., N'debi,M., Demontant,V., Trawinski,E.
- EPI_ISL_447656; hCoV-19/France/40004SR/2020; 2020-05-12; Europe / France; Genomic platform; De Prost,N., Fourati,S., Lamoureux,C., Schmitz,D., Deveaux,I., Picard,O., Lepeule,R., Surgers,L., Mekontso-Dessap,A., Woerther,P.-L., Canoui-Poitrine,F., Pawlotsky,J.-M., Clinical Study Group,C., Rodrigue,C., Gricourt,G., N'debi,M., Demontant,V., Trawinski,E.

- EPI_ISL_447692; hCoV-19/France/40007RD/2020; 2020-05-14; Europe / France; Genomic platform; De Prost,N., Fourati,S., Lamoureux,C., Schmitz,D., Deveaux,I., Picard,O., Lepeule,R., Surgers,L., Mekontso-Dessap,A., Woerther,P.-L., Canoui-Poitrine,F., Pawlotsky,J.-M., Clinical Study Group,C., Rodrigue,C., Gricourt,G., N'debi,M., Demontant,V., Trawinski,E.
- EPI_ISL_447694; hCoV-19/France/50002BS/2020; 2020-05-14; Europe / France; Genomic platform; De Prost,N., Fourati,S., Lamoureux,C., Schmitz,D., Deveaux,I., Picard,O., Lepeule,R., Surgers,L., Mekontso-Dessap,A., Woerther,P.-L., Canoui-Poitrine,F., Pawlotsky,J.-M., Clinical Study Group,C., Rodrigue,C., Gricourt,G., N'debi,M., Demontant,V., Trawinski,E.
- EPI_ISL_447716; hCoV-19/France/50011LM/2020; 2020-05-14; Europe / France; Genomic platform; De Prost,N., Fourati,S., Lamoureux,C., Schmitz,D., Deveaux,I., Picard,O., Lepeule,R., Surgers,L., Mekontso-Dessap,A., Woerther,P.-L., Canoui-Poitrine,F., Pawlotsky,J.-M., Clinical Study Group,C., Rodrigue,C., Gricourt,G., N'debi,M., Demontant,V., Trawinski,E.
- EPI_ISL_420039; hCoV-19/France/GE2720/2020; 2020-04-04; Europe / France / Grand-Est / Strasbourg; National Reference Center for Viruses of Respiratory Infections, Institut Pasteur, Paris; Mélanie Albert, Marion Barbet, Sylvie Behillil, Méline Bizard, Angela Brisebarre, Flora Donati, Etienne Simon-Lorière, Vincent Enouf, Maud Vanpeene, Sylvie van der Werf.
- EPI_ISL_420040; hCoV-19/France/GE2722/2020; 2020-04-04; Europe / France / Grand-Est / Strasbourg; National Reference Center for Viruses of Respiratory Infections, Institut Pasteur, Paris; Mélanie Albert, Marion Barbet, Sylvie Behillil, Méline Bizard, Angela Brisebarre, Flora Donati, Etienne Simon-Lorière, Vincent Enouf, Maud Vanpeene, Sylvie van der Werf.
- EPI_ISL_420044; hCoV-19/France/HF2797/2020; 2020-04-04; Europe / France / Hauts de France / Chateau-Thierry; National Reference Center for Viruses of Respiratory Infections, Institut Pasteur, Paris; Mélanie Albert, Marion Barbet, Sylvie Behillil, Méline Bizard, Angela Brisebarre, Flora Donati, Etienne Simon-Lorière, Vincent Enouf, Maud Vanpeene, Sylvie van der Werf.
- EPI_ISL_418240; hCoV-19/France/IDF2684/2020; 2020-03-29; Europe / France / Ile de France / Longjumeau; National Reference Center for Viruses of Respiratory Infections, Institut Pasteur, Paris; Mélanie Albert, Marion Barbet, Sylvie Behillil, Méline Bizard, Angela Brisebarre, Flora Donati, Etienne Simon-Lorière, Vincent Enouf, Maud Vanpeene, Sylvie van der Werf.
- EPI_ISL_421512; hCoV-19/France/IDF3324/2020; 2020-04-08; Europe / France / Ile de France / Longjumeau; National Reference Center for Viruses of Respiratory Infections, Institut Pasteur, Paris; Mélanie Albert, Marion Barbet, Sylvie Behillil, Méline Bizard, Angela Brisebarre, Flora Donati, Etienne Simon-Lorière, Vincent Enouf, Maud Vanpeene, Sylvie van der Werf.
- EPI_ISL_428352; hCoV-19/France/IDF3518/2020; 2020-04-20; Europe / France / Ile De France / Orsay; National Reference Center for Viruses of Respiratory Infections, Institut Pasteur, Paris; Mélanie Albert, Marion Barbet, Sylvie Behillil, Méline Bizard, Angela Brisebarre, Flora Donati, Etienne Simon-Lorière, Vincent Enouf, Maud Vanpeene, Sylvie van der Werf.
- EPI_ISL_444969; hCoV-19/Guangdong/SYSU-IHV/2020; 2020-05-14; Asia / China / Guangdong / Guangzhou; Institute of Human Virology, Zhongshan School of Medicine, Sun Yat-sen University; Junsong Zhang, Fei Yu, Jun Liu, Huimin Fan, Ruosu Ying, Feng Huang, Ting Pan, Bingfeng Liu,Yiwen Zhang, Xu Zhang, Mang Shi, Fengyu Hu, Fang Li, Kai Deng, Hui Zhang.
- EPI_ISL_421746; hCoV-19/Luxembourg/LNS0819394/2020; 2020-04-09; Europe / Luxembourg; Laboratoire National de Sante, Microbiology, Epidemiology and Microbial Genomics; Anke Wienecke-Baldacchino, Ardashel Latsuzbaia, Jessica Tapp, Catherine Ragimbeau, Guillaume Fournier, Tamir Abdelrahman, Trung Nguyen Nguyen, Joel Mossong.
- EPI_ISL_434493; hCoV-19/Luxembourg/LNS1368952/2020; 2020-04-29; Europe / Luxembourg; Laboratoire National de Sante, Microbiology, Epidemiology and Microbial Genomics; Anke Wienecke-Baldacchino, Ardashel Latsuzbaia, Jessica Tapp, Catherine Ragimbeau, Guillaume Fournier, Tamir Abdelrahman, Trung Nguyen Nguyen, Joel Mossong.
- EPI_ISL_445069; hCoV-19/Luxembourg/LNS3029697/2020; 2020-05-14; Europe / Luxembourg; Laboratoire National de Sante, Microbiology, Epidemiology and Microbial Genomics; Anke Wienecke-Baldacchino, Ardashel Latsuzbaia, Jessica Tapp, Catherine Ragimbeau, Guillaume Fournier, Tamir Abdelrahman, Trung Nguyen Nguyen, Joel Mossong.
- EPI_ISL_421752; hCoV-19/Luxembourg/LNS4569788/2020; 2020-04-09; Europe / Luxembourg; Laboratoire National de Sante, Microbiology, Epidemiology and Microbial Genomics; Anke Wienecke-Baldacchino, Ardashel Latsuzbaia, Jessica Tapp, Catherine Ragimbeau, Guillaume Fournier, Tamir Abdelrahman, Trung Nguyen Nguyen, Joel Mossong.

- EPI_ISL_429208; hCoV-19/Switzerland/GE2164/2020; 2020-04-23; Europe / Switzerland; University Hospitals of Geneva Laboratory of Virology; Laubscher F.
- EPI_ISL_463893; hCoV-19/Bosnia and Herzegovina/03_Tuzla/2020; 2020-06-10; Europe / Bosnia and Herzegovina / Tuzla; Alea Genetički Centar; Konjhodžić,R;Salihefendić,L;Goletić,T;Pećar,D;Tihić,N;Marjanović,D;Hukić,M.
- EPI_ISL_462455; hCoV-19/Bosnia and Herzegovina/ChVir7348/2020; 2020-06-08; Europe / Bosnia and Herzegovina / Sarajevo; Charite Universitatsmedizin Berlin, Institute of Virology; Victor M Corman, Jorn Beheim-Schwarzbach, Barbara Muehlemann, Talitha Veith, Julia Schneider, Terry Jones, Amela Dedeic-Ljubovic, Irma Salimovic-Besic, Suzana Arapcic, Almedina Hadzihasanovic-Moro, Selma Mutevelic, Christian Drosten.
- EPI_ISL_462456; hCoV-19/Bosnia and Herzegovina/ChVir7349/2020; 2020-06-08; Europe / Bosnia and Herzegovina / Sarajevo; Charite Universitatsmedizin Berlin, Institute of Virology; Victor M Corman, Jorn Beheim-Schwarzbach, Barbara Muehlemann, Talitha Veith, Julia Schneider, Terry Jones, Amela Dedeic-Ljubovic, Irma Salimovic-Besic, Suzana Arapcic, Almedina Hadzihasanovic-Moro, Selma Mutevelic, Christian Drosten.
- EPI_ISL_462457; hCoV-19/Bosnia and Herzegovina/ChVir7350/2020; 2020-06-08; Europe / Bosnia and Herzegovina / Sarajevo; Charite Universitatsmedizin Berlin, Institute of Virology; Victor M Corman, Jorn Beheim-Schwarzbach, Barbara Muehlemann, Talitha Veith, Julia Schneider, Terry Jones, Amela Dedeic-Ljubovic, Irma Salimovic-Besic, Suzana Arapcic, Almedina Hadzihasanovic-Moro, Selma Mutevelic, Christian Drosten.
- EPI_ISL_462465; hCoV-19/Bosnia and Herzegovina/ChVir7362/2020; 2020-06-08; Europe / Bosnia and Herzegovina / Sarajevo; Charite Universitatsmedizin Berlin, Institute of Virology; Victor M Corman, Jorn Beheim-Schwarzbach, Barbara Muehlemann, Talitha Veith, Julia Schneider, Terry Jones, Amela Dedeic-Ljubovic, Irma Salimovic-Besic, Suzana Arapcic, Almedina Hadzihasanovic-Moro, Selma Mutevelic, Christian Drosten.
- EPI_ISL_462469; hCoV-19/Bosnia and Herzegovina/ChVir7367/2020; 2020-06-08; Europe / Bosnia and Herzegovina / Sarajevo; Charite Universitatsmedizin Berlin, Institute of Virology; Victor M Corman, Jorn Beheim-Schwarzbach, Barbara Muehlemann, Talitha Veith, Julia Schneider, Terry Jones, Amela Dedeic-Ljubovic, Irma Salimovic-Besic, Suzana Arapcic, Almedina Hadzihasanovic-Moro, Selma Mutevelic, Christian Drosten.
- EPI_ISL_448540; hCoV-19/England/OXON-ACE2A/2020; 2020-05-19; Europe / United Kingdom / England; COVID-19 Genomics UK (COG-UK) Consortium; Tanya Golubchik, David Bonsall, George Macintyre, Amy Trebes, Mariateresa de Cesare, Catrin Moore, Alex Mobbs, Anita Justice, Robert Shaw, Monique Andersson, Emma Wise, Nathan Moore, Jessica Lynch, Nick Cortes, Stephen Kidd, David Buck, John Todd, Christophe Fraser.
- EPI_ISL_448797; hCoV-19/England/OXON-AEF43/2020; 2020-05-19; Europe / United Kingdom / England; COVID-19 Genomics UK (COG-UK) Consortium; Tanya Golubchik, David Bonsall, George Macintyre, Amy Trebes, Mariateresa de Cesare, Catrin Moore, Alex Mobbs, Anita Justice, Robert Shaw, Monique Andersson, Emma Wise, Nathan Moore, Jessica Lynch, Nick Cortes, Stephen Kidd, David Buck, John Todd, Christophe Fraser.
- EPI_ISL_457727; hCoV-19/Estonia/ChVir4287/2020; 2020-06-02; Europe / Estonia; Charite Universitatsmedizin Berlin, Institute of Virology; Victor M Corman, Jorn Beheim-Schwarzbach, Barbara Mühlemann, Talitha Veith, Julia Schneider, Paul Naaber, Terry Jones, Christian Drosten.
- EPI_ISL_455643; hCoV-19/India/S13/2020; 2020-06-01; Asia / India / West_Bengal / North 24 Parganas; National Institute of Biomedical Genomics; Arindam Maitra, Mamta Chawla Sarkar, Sreedhar Chinnaswamy, Hasina Banu, Ananya Chatterjee, Shanta Dutta, Saumitra Das.
- EPI_ISL_457700; hCoV-19/Italy/2629/2020; 2020-06-02; Europe / Italy / PA Trento; Army Medical and Veterinary Research Center; Paola Stefanelli, Alessandra Lo Presti, Stefano Fiore, Antonella Marchi, Eleonora Benedetti, Concetta Fabiani Silvia Fillo, Giovanni Faggioni, Riccardo De Sanctis, Antonella Fortunato, Anna Anselmo, Francesco Giordani, Vanessa Vera Fain, Nino D'Amore, Florigio Lista.
- EPI_ISL_451308; hCoV-19/Italy/LO-13075-B/2020; 2020-05-24; Europe / Italy / Lombardia; Laboratory of Virology, INMI Lazzaro Spallanzani IRCCS; Antonio Piralla, Fausto Baldanti, Maria R. Capobianchi, Cesare E.M. Gruber, Martina Rueca, Barbara Bartolini, Antonino Di Caro.

- EPI_ISL_451309; hCoV-19/Italy/LO-13075-N/2020; 2020-05-24; Europe / Italy / Lombardia; Laboratory of Virology, INMI Lazzaro Spallanzani IRCCS; Fausto Baldanti, Antonio Piralla, Cesare E.M. Gruber, Maria R. Capobianchi, Antonino Di Caro, Martina Rueca, Barbara Bartolini.
- EPI_ISL_451306; hCoV-19/Italy/PV-5314-B/2020; 2020-05-24; Europe / Italy / Lombardia; Laboratory of Virology, INMI Lazzaro Spallanzani IRCCS; Antonio Piralla, Fausto Baldanti, Martina Rueca, Antonino Di Caro, Maria R. Capobianchi, Cesare E.M. Gruber, Barbara Bartolini.
- EPI_ISL_455444; hCoV-19/Poland/PL_P31/2020; 2020-05-29; Europe / Poland / Warminsko-Mazurskie / Olsztyn; 1. ViroGenetics - BSL3 Laboratory of Virology, Małopolska Centre of Biotechnology, Jagiellonian University; 2. II Department of Internal Medicine, Faculty of Medicine, Jagiellonian University Medical College; 3. Narodowy Instytut Zdrowia Publicznego – Państwowy Zakład Higieny (NIZP-PZH); Katarzyna Pancer, Marek Sanak, Aleksandra A. Zasada, Magdalena Rzeczkowska, Tomasz Wołkowicz, Katarzyna Zacharczuk, Agnieszka Kołakowska-Kulesza, Katarzyna Owczarek, Aleksandra Milewska, Natalia Wolaniuk, Ewelina Hallman-Szelińska, Paweł P Łabaj, Wojciech Branicki, Krzysztof Pyrć.
- EPI_ISL_437754; hCoV-19/Saudi Arabia/KAUST-Jeddah61/2020; 2020-05-11; Asia / Saudi Arabia / Jeddah; Pathogen Genomics Lab King Abdullah University of Science and Technology(KAUST); Sharif Hala,Fadwa Alofi,Afrah Alsomali, Asim Khogeer, Sara Mfarrej, Khaled Algithami,Raeece Naeem, Amit Kumar Subudhi,Fathia Ben-Rached, Rahul Salunke, Anwar Hashem, Naif Almontashiri, Arnab Pain.
- EPI_ISL_454456; hCoV-19/Sweden/20-08082/2020; 2020-05-28; Europe / Sweden / Stockholm; The Public Health Agency of Sweden; Anna-Malin Linde, Maria Lind Karlberg, Mattias Haukland, Reza Advani, Olov Svartstrom, Oskar Karlsson Lindsjo, Petra Edquist, Shamam Muradrasoli, Anna Risberg, Karin Tegmark-Wisell.
- EPI_ISL_454475; hCoV-19/Sweden/20-08104/2020; 2020-05-28; Europe / Sweden / Stockholm; The Public Health Agency of Sweden; Anna-Malin Linde, Maria Lind Karlberg, Mattias Haukland, Reza Advani, Olov Svartstrom, Oskar Karlsson Lindsjo, Petra Edquist, Shamam Muradrasoli, Anna Risberg, Karin Tegmark-Wisell.
- EPI_ISL_454487; hCoV-19/Sweden/20-08116/2020; 2020-05-28; Europe / Sweden / Stockholm; The Public Health Agency of Sweden; Anna-Malin Linde, Maria Lind Karlberg, Mattias Haukland, Reza Advani, Olov Svartstrom, Oskar Karlsson Lindsjo, Petra Edquist, Shamam Muradrasoli, Anna Risberg, Karin Tegmark-Wisell.
- EPI_ISL_455847; hCoV-19/Sweden/20-08117/2020; 2020-06-01; Europe / Sweden / Stockholm; The Public Health Agency of Sweden; Anna-Malin Linde, Maria Lind Karlberg, Mattias Haukland, Reza Advani, Olov Svartstrom, Oskar Karlsson Lindsjo, Petra Edquist, Shamam Muradrasoli, Anna Risberg, Karin Tegmark-Wisell.
- EPI_ISL_454880; hCoV-19/Sweden/20-50104/2020; 2020-05-29; Europe / Sweden / Stockholm; The Public Health Agency of Sweden; Anna-Malin Linde, Maria Lind Karlberg, Mattias Haukland, Reza Advani, Olov Svartstrom, Oskar Karlsson Lindsjo, Petra Edquist, Shamam Muradrasoli, Anna Risberg, Karin Tegmark-Wisell.
- EPI_ISL_455876; hCoV-19/Sweden/20-50219/2020; 2020-06-01; Europe / Sweden / Stockholm; The Public Health Agency of Sweden; Anna-Malin Linde, Maria Lind Karlberg, Mattias Haukland, Reza Advani, Olov Svartstrom, Oskar Karlsson Lindsjo, Petra Edquist, Shamam Muradrasoli, Anna Risberg, Karin Tegmark-Wisell.
- EPI_ISL_455878; hCoV-19/Sweden/20-50223/2020; 2020-06-01; Europe / Sweden / Stockholm; The Public Health Agency of Sweden; Anna-Malin Linde, Maria Lind Karlberg, Mattias Haukland, Reza Advani, Olov Svartstrom, Oskar Karlsson Lindsjo, Petra Edquist, Shamam Muradrasoli, Anna Risberg, Karin Tegmark-Wisell.
- EPI_ISL_451197; hCoV-19/Uganda/UG015/2020; 2020-05-23; Africa / Uganda; MRC/UVRI & LSHTM Uganda Research Unit; Dan Lule Bugembe, John Kayiwa, My V.T Phan, Phiona Tushabe, Stephen Balinandi, Beatrice Dhaala, Deogratius Ssemwanga, Jonas Lexow, Henry Mwebesa, Jane Aceng, Henry Kyobe, Julius Lutwama, Pontiano Kaleebu, Matthew Cotton.
- EPI_ISL_454637; hCoV-19/USA/CA-CZB-1279/2020; 2020-05-28; North America / USA / California / Humboldt County; Chan-Zuckerberg Biohub; CZB Cliahub Consortium.
- EPI_ISL_447840; hCoV-19/USA/DC-CDC-0019/2020; 2020-05-18; North America / USA / Washington DC; Pathogen Discovery, Respiratory Viruses Branch, Division of Viral Diseases, Centers for Disease Control and Prevention; Krista Queen, Yan Li, Anna Uehara, Jing Zhang,

Ying Tao, Clinton R. Paden, Haibin Wang, Jasmine Padilla, Mary S. Keckler, Alison S. Laufer Halpin, Justin Lee, Christopher A. Elkins, Suxiang Tong.

- EPI_ISL_460471; hCoV-19/USA/MA-MGH-00205/2020; 2020-05-26; North America / USA / Massachusetts; Infectious Disease Program, Broad Institute of Harvard and MIT; Lemieux,J.E., Siddle,K.J., Shaw,B., Adams,G., Pierce,V., Turbett,S., Anahtar,M., Branda,J., Slater,D., Harris,J., Lin,A.E., Gladden-Young,A., Lagerborg,K., Rudy,M., DeRuff,K., Carter,A., Normandin,E., Bauer,M., Reilly,S., Tomkins-Tinch,C., Loreth,C., Chaluvadi,S., Neumann,A., Cusick,C., Chapman,S.B., Gnirke,A., Flowers,K., Cerrato,F., Birren,B.W., Gallagher,G., Smole,S., Park,D.J., MacInnis,B.L., Ryan,E., LaRocque,R., Rosenberg,E., Sabeti,P.C.
- EPI_ISL_444656; hCoV-19/USA/NY-NYUMC539/2020; 2020-05-14; North America / USA / New York / Queens; Departments of Pathology and Medicine, New York University School of Medicine; Maria Aguero-Rosenfeld, Brendan Belovarac, Margaret Black, Ludovic Boytard, John Cadley, Paolo Cotzia, John Chen, Dacia Dimartino, Xiaojun Feng, Tatyana Gindin, Emily Guzman, Adriana Heguy, Megan Hogan, Emily Huang, George Jour, Alireza Khodadadi-Jamayran, Lawrence H. Lin, Raven Luther, Andrew Lytle, Christian Marier, Matthew T. Maurano, Mark J. Mulligan, Peter Meyn, Raquel Ordonez Ciriza, Iman Osman, Jared Pinnell, Vanessa Raabe, Sitharam Ramaswami, Amy Rapkiewicz, Andre M. Ribeiro-dos-Santos, Marie Samanovic-Golden, Antonio Serrano, Guomiao Shen, Matija Snuderl, Theodore Vougiouklakis, Nick Vulpescu, Gael Westby, Paul Zappile, Yutong Zhang.
- EPI_ISL_436468; hCoV-19/USA/PA-MGSC13-04/2020; ; 2020-05-06; North America / USA / Pennsylvania / Pittsburgh; Microbial Genome Sequencing Center, Microbial Genomic Epidemiological Laboratory; Dan Snyder, Stephanie L Mitchell, Mustapha M Mustapha, Marissa P Griffith, Vatsala R Srinivasa, Kady D Waggle, Chinelo Ezeonwuku, Jane W. Marsh, Lee H. Harrison, Vaughn S. Cooper.

The authors gratefully acknowledge the Authors and Submitting laboratories for sharing their sequence in NCBI database. Its GenBank Accession ID, name, date of submission and authors are displayed below:

- MN908947.3; Severe acute respiratory syndrome coronavirus 2 isolate Wuhan-Hu-1; 2020-01-05; Wu,F., Zhao,S., Yu,B., Chen,Y.M., Wang,W., Song,Z.G., Hu,Y., Tao,Z.W., Tian,J.H., Pei,Y.Y., Yuan,M.L., Zhang,Y.L., Dai,F.H., Liu,Y., Wang,Q.M., Zheng,J.J., Xu,L., Holmes,E.C. and Zhang,Y.Z.

**Figure S2. Maximum likelihood uncondensed phylogenetic tree (below).**

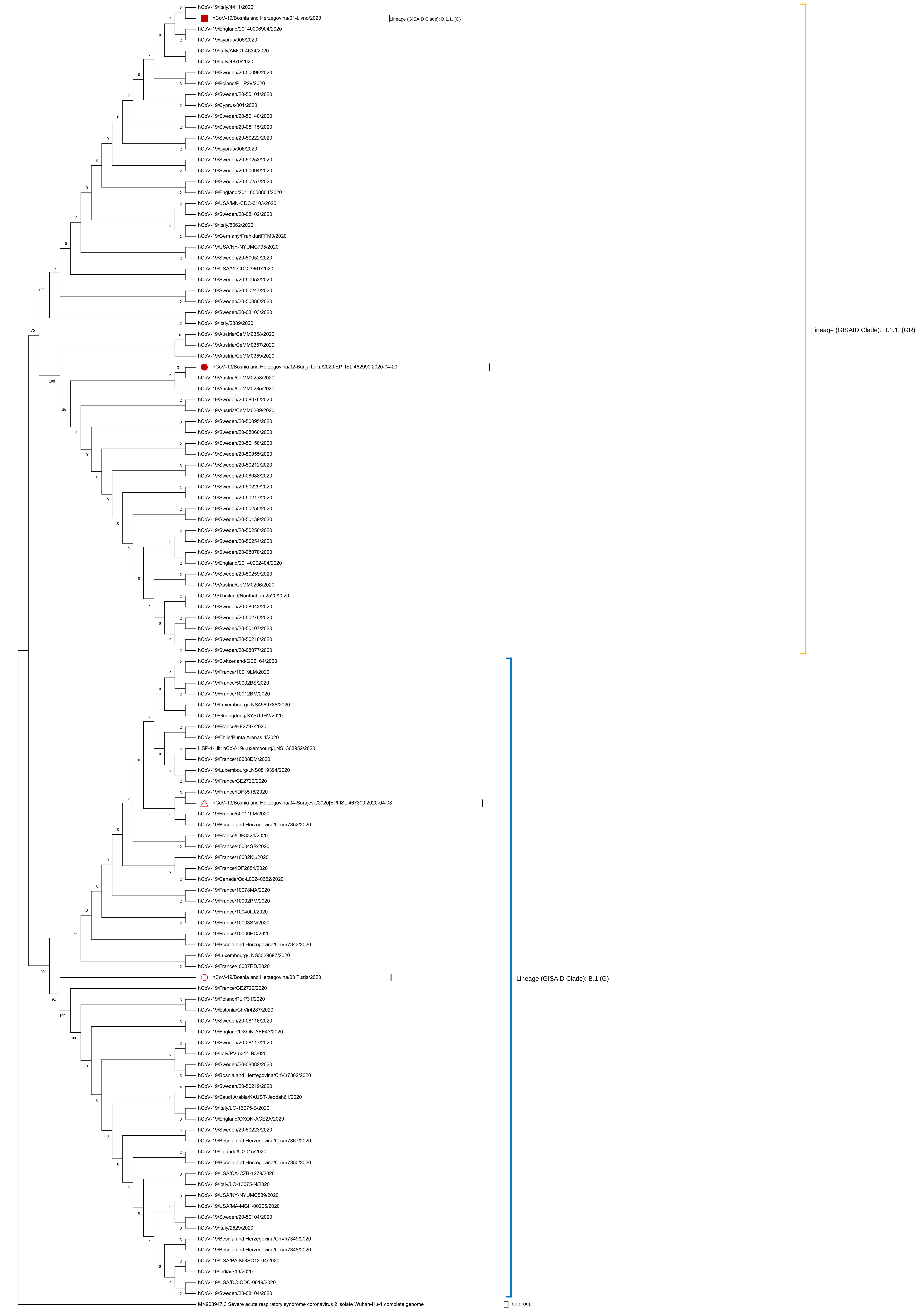
hCoV-19/Italy/4411/2020
hCoV-19/Bosnia and Herzegovina/01-Livno/2020
Lineage (GISAID Clade): B.1.1. (O)
hCoV-19/England/20140006904/2020
hCoV-19/Cyprus/005/2020
hCoV-19/Italy/AMC1-4634/2020
hCoV-19/Italy/4870/2020
hCoV-19/Sweden/20-50098/2020
hCoV-19/Poland/PL P29/2020
hCoV-19/Sweden/20-50101/2020
hCoV-19/Cyprus/001/2020
hCoV-19/Sweden/20-50140/2020
hCoV-19/Sweden/20-08115/2020
hCoV-19/Sweden/20-50222/2020
hCoV-19/Cyprus/006/2020
hCoV-19/Sweden/20-50253/2020
hCoV-19/Sweden/20-50094/2020
hCoV-19/Sweden/20-50257/2020
hCoV-19/England/20118050804/2020
hCoV-19/USA/MN-CDC-0103/2020
hCoV-19/Sweden/20-08102/2020
hCoV-19/Italy/5062/2020
hCoV-19/Germany/FrankfurtFFM3/2020
hCoV-19/USA/NY-NYUMC795/2020
hCoV-19/Sweden/20-50052/2020
hCoV-19/USA/VI-CDC-3661/2020
hCoV-19/Sweden/20-50053/2020
hCoV-19/Sweden/20-50247/2020
hCoV-19/Sweden/20-50088/2020
hCoV-19/Sweden/20-08103/2020
hCoV-19/Italy/2389/2020
hCoV-19/Austria/CeMM0358/2020
hCoV-19/Austria/CeMM0357/2020
hCoV-19/Austria/CeMM0359/2020
hCoV-19/Bosnia and Herzegovina/02-Banja Luka/2020|EPI ISL 462990|2020-04-29
hCoV-19/Austria/CeMM0258/2020
hCoV-19/Austria/CeMM0285/2020
hCoV-19/Sweden/20-08076/2020
hCoV-19/Austria/CeMM0209/2020
hCoV-19/Sweden/20-50095/2020
hCoV-19/Sweden/20-08060/2020
hCoV-19/Sweden/20-50150/2020
hCoV-19/Sweden/20-50055/2020
hCoV-19/Sweden/20-50212/2020
hCoV-19/Sweden/20-08088/2020
hCoV-19/Sweden/20-50229/2020
hCoV-19/Sweden/20-50217/2020
hCoV-19/Sweden/20-50255/2020
hCoV-19/Sweden/20-50139/2020
hCoV-19/Sweden/20-50256/2020
hCoV-19/Sweden/20-50254/2020
hCoV-19/Sweden/20-08078/2020
hCoV-19/England/20140002404/2020
hCoV-19/Sweden/20-50259/2020
hCoV-19/Austria/CeMM0206/2020
hCoV-19/Thailand/Nonthaburi 2520/2020
hCoV-19/Sweden/20-08043/2020
hCoV-19/Sweden/20-50270/2020
hCoV-19/Sweden/20-50107/2020
hCoV-19/Sweden/20-50218/2020
hCoV-19/Sweden/20-08077/2020
Lineage (GISAID Clade): B.1.1. (GR)
hCoV-19/Switzerland/GE2164/2020
hCoV-19/France/10019LM/2020
hCoV-19/France/50002BS/2020
hCoV-19/France/10012BM/2020
hCoV-19/Luxembourg/LNS4569788/2020
hCoV-19/Guangdong/SYSU-IHV/2020
hCoV-19/France/HF2797/2020
hCoV-19/Chile/Punta Arenas 4/2020
HSP-1-Hit: hCoV-19/Luxembourg/LNS1368952/2020
hCoV-19/France/10008DM/2020
hCoV-19/Luxembourg/LNS0819394/2020
hCoV-19/France/GE2720/2020
hCoV-19/France/IDF3518/2020
hCoV-19/Bosnia and Herzegovina/04-Sarajevo/2020|EPI ISL 467300|2020-04-08
hCoV-19/France/50011LM/2020
hCoV-19/Bosnia and Herzegovina/ChVir7352/2020
hCoV-19/France/IDF3324/2020
hCoV-19/France/40004SR/2020
hCoV-19/France/10032KL/2020
hCoV-19/France/IDF2684/2020
hCoV-19/Canada/Qc-L00240652/2020
hCoV-19/France/10078MA/2020
hCoV-19/France/10002PM/2020
hCoV-19/France/10040LJ/2020
hCoV-19/France/10003SN/2020
hCoV-19/France/10006HC/2020
hCoV-19/Bosnia and Herzegovina/ChVir7343/2020
hCoV-19/Luxembourg/LNS3029697/2020
hCoV-19/France/40007RD/2020
hCoV-19/Bosnia and Herzegovina/03 Tuzla/2020
Lineage (GISAID Clade): B.1 (G)
hCoV-19/France/GE2722/2020
hCoV-19/Poland/PL P31/2020
hCoV-19/Estonia/ChVir4287/2020
hCoV-19/Sweden/20-08116/2020
hCoV-19/England/OXON-AEF43/2020
hCoV-19/Sweden/20-08117/2020
hCoV-19/Italy/PV-5314-B/2020
hCoV-19/Sweden/20-08082/2020
hCoV-19/Bosnia and Herzegovina/ChVir7362/2020
hCoV-19/Sweden/20-50219/2020
hCoV-19/Saudi Arabia/KAUST-Jeddah61/2020
hCoV-19/Italy/LO-13075-B/2020
hCoV-19/England/OXON-ACE2A/2020
hCoV-19/Sweden/20-50223/2020
hCoV-19/Bosnia and Herzegovina/ChVir7367/2020
hCoV-19/Uganda/UG015/2020
hCoV-19/Bosnia and Herzegovina/ChVir7350/2020
hCoV-19/USA/CA-CZB-1279/2020
hCoV-19/Italy/LO-13075-N/2020
hCoV-19/USA/NY-NYUMC539/2020
hCoV-19/USA/MA-MGH-00205/2020
hCoV-19/Sweden/20-50104/2020
hCoV-19/Italy/2629/2020
hCoV-19/Bosnia and Herzegovina/ChVir7349/2020
hCoV-19/Bosnia and Herzegovina/ChVir7348/2020
hCoV-19/USA/PA-MGSC13-04/2020
hCoV-19/India/S13/2020
hCoV-19/USA/DC-CDC-0019/2020
hCoV-19/Sweden/20-08104/2020
MN908947.3 Severe acute respiratory syndrome coronavirus 2 isolate Wuhan-Hu-1 complete genome
outgroup
